# Novel estimators for family-based genome-wide association studies increase power and robustness

**DOI:** 10.1101/2022.10.24.513611

**Authors:** Junming Guan, Seyed Moeen Nehzati, Daniel J. Benjamin, Alexander I. Young

## Abstract

A goal of genome-wide association studies (GWASs) is to estimate the causal effects of alleles carried by an individual on that individual (‘direct genetic effects’). Typical GWAS designs, however, are susceptible to confounding due to gene-environment correlation and non-random mating (population stratification and assortative mating). Family-based GWAS, in contrast, is robust to such confounding since it uses random, within-family genetic variation. When both parents are genotyped, a regression controlling for parental genotype provides the most powerful approach. However, parental genotypes are often missing. We have previously shown that imputing the genotypes of missing parent(s) can increase power for estimation of direct genetic effects over using genetic differences between siblings. We extend the imputation method, which previously only applied to samples with at least one genotyped sibling or parent, to ‘singletons’ (individuals without any genotyped relatives). By including singletons, the effective sample size for estimation of direct effects can be increased by up to 50%. We apply this method to 408,254 ‘White British’ individuals from the UK Biobank, obtaining an effective sample size increase of between 25% and 43% (depending upon phenotype) by including 368,629 singletons. While this approach maximizes power, it can be biased when there is strong population structure. We therefore introduce an imputation based estimator that is robust to population structure and more powerful than other robust estimators. We implement our estimators in the software package snipar using an efficient linear-mixed model (LMM) specified by a sparse genetic relatedness matrix. We examine the bias and variance of different family-based and standard GWAS estimators theoretically and in simulations with differing levels of population structure, enabling researchers to choose the appropriate approach depending on their research goals.

## 1 Introduction

Genome-wide association studies (GWASs) have identified thousands of associations between genetic variants and human phenotypes and diseases[1]. Standard GWAS study designs estimate the association between a phenotype and an allele by regression of individuals’ phenotypes onto the number of copies of alleles they carry. Multiple phenomena contribute to the associations estimated by standard GWAS, which we call ‘population effects’, as they reflect the genotype-phenotype association in the population[2]: causal effects of alleles carried by the individual on the individual (direct genetic effects); effects of alleles in relative(s) through the environment, called indirect genetic effects (IGEs) or genetic nurture[3]; and confounding due to population stratification and assortative mating (AM)[4, 5], which result in correlations between the genetic variant and other genetic and environmental factors[3, 6–8]. Adjustment for genetic principal components and linear mixed models (LMMs) reduce confounding due to population stratification[4, 5] and AM [9], but studies have shown that this often fails to remove all confounding[5, 6, 8, 9].

The only known methods that are guaranteed to remove confounding due to IGEs and non-random mating are family-based methods, such as family-based GWAS[8, 10]. The family-based GWAS regression[8] adds parental genotype as a ‘control’:

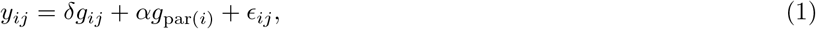

where *y_ij_* is the phenotype of the *j*th sibling of the *i*th family; *g_ij_* is the corresponding genotype; *g*_par(*i*)_ = *g*_*p*(*i*)_ + *g*_*m*(*i*)_ is the sum of paternal and maternal genotypes; *δ* is the direct genetic effect; *α* is the average non-transmitted coefficient (NTC); and *ϵ_ij_* is the residual. Since 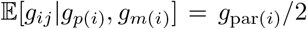, and variation in offspring genotype around this expectation is due to random Mendelian segregations, estimates of direct genetic effects from fitting Model 1 are free from confounding due to IGEs and non-random mating[8]. The average NTC captures IGEs from relatives and confounding due to population stratification and assortative mating[8]. Alternatively, one can fit a model that allows for different coefficients on the paternal and maternal genotypes:

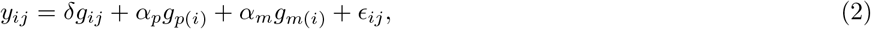

where *α_p_* and *α_m_* are, respectively, the paternal and maternal NTCs. Model 1 can be derived from Model 2 (with a change of residuals)[8], implying that *α* = (*α_p_* + *α_m_*)/2. Although Model 1 is sufficient to remove confounding from estimates of direct effects, irrespective of whether *α_p_* = *α_m_*, Model 2 may be preferred in certain contexts (Appendix A), such as if one is interested in differences between *α_p_* and *α_m_*. Standard GWAS performs a regression of phenotype onto genotype, giving an estimate of the population effect, *β*. Assuming random-mating, it can be shown that[8]: *β* = *δ* + *α*. This provides a useful connection between the parameters of family-based and standard GWAS.

Fitting Models 1 and 2 entails restricting one’s sample to those with both parents genotyped, which is often only a small fraction (or none) of the available samples. An alternative is to analyze genetic differences between siblings[9, 10]. For example, one can perform a regression of phenotype differences between siblings onto genotype differences between siblings:

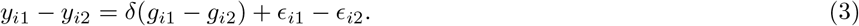

Estimates of *δ* from sibling differences are free from confounding due to non-random mating and IGEs, except for IGEs from siblings[8]. However, provided that one models the correlation between siblings’ residuals (as in a generalized least-squares estimator[8]), estimates of direct genetic effects from Models 1 and 2 are more precise than from genetic differences between siblings. Furthermore, using genetic differences between siblings results in ignoring samples with genotyped parent(s) but without genotyped siblings[10].

Young et al. [8] proposed an alternative approach: to treat parental genotypes as missing data and to impute them according to Mendelian laws. Consider the case of a sibling pair. Young et al. [8] propose to impute the missing parental genotype, *g*_par(*i*)_, conditional on the identical-by-descent (IBD) state of the siblings; i.e., whether the siblings have inherited the same or different alleles from each parent. For sibling pairs that do not share any alleles IBD (IBD0), all four parental alleles have been observed, so the imputed parental genotype is the sum of sibling genotypes; for sibling pairs that share one allele IBD (IBD1), three parental alleles have been observed, and the imputed parental genotype is the sum of the three parental alleles plus the population allele frequency, *f*, which is used in place of the unobserved parental allele; and for sibling pairs that share both alleles IBD (IBD2), only two parental alleles have been observed, and the imputed parental genotype is *g*_*i*1_ + 2*f* = *g*_*i*2_ + 2*f*. On average, 3 parental alleles can be recovered by imputation, and the resulting imputed parental genotypes are not a linear function of sibling genotypes across the different IBD states. The imputed parental genotypes are then used in place of the observed ones to perform the regressions given by Models 1 and 2. Provided that the imputed parental genotypes are unbiased estimates of the true parental genotypes, the resulting direct effect estimates are unbiased and consistent, and the empirical sampling variance-covariance matrix is an unbiased estimate of the true sampling variance-covariance matrix[8]. This approach increases the effective sample size for direct genetic effects by up to 1/3rd compared to using genetic differences between siblings. Beyond genotyped sibling pairs, the imputation method of Young et al. enables the inclusion of any genotyped sample with at least one genotyped first-degree relative, including samples with one or both parent(s) genotyped but without genotyped siblings, further increasing power[8].

However, the method of Young et al. ignores most of the genotyped sample in large-scale biobanks such as the UK Biobank, where only ~10% of the sample have a genotyped first-degree relative[11]. Large samples of individuals without genotyped relatives, referred to here as ‘singletons’, can provide precise estimates of *β*, the population effect (as in standard GWAS). Since, under random-mating[8], *β* = *δ* + *α* = *δ* + (*α_p_* + *α_m_*)/2, a precise estimate of the population effect puts a constraint on the set of plausible values the family-based GWAS parameter vector *θ* = (*δ, α_p_, α_m_*) can take. Following this intuition, we show that, under random-mating, including singletons by linearly imputing their parents’ genotypes can increase the effective sample size for direct effects by up to 50% over the method of Young et al. Furthermore, by including singletons, we are able to estimate the population effect with similar precision to standard GWAS, thereby creating a ‘unified’ estimator for both direct and population effects. We apply this method to 408,254 ‘White British’ individuals from the UK Biobank (UKB), obtaining an effective sample size increase for direct effects of between 25% and 43% (across 20 phenotypes) by including 368,629 singletons.

While the unified estimator maximizes power for estimation of direct genetic effects, we show that population structure can cause bias in the imputed parental genotypes and downstream analysis. We therefore develop an imputation-based estimator that is robust to population structure, which we call the ‘robust estimator’, and is more powerful than alternative approaches that are robust to population structure.

We implement the different estimators in a linear mixed model in *snipar*. The linear mixed model includes a random effect for differences in phenotypic mean between sibships (as in Young et al.[8]) and a random effect specified by a sparse genetic relatedness matrix (GRM), as in *fastGWA*[12]. We examine the family-based and standard GWAS estimators in a bias-variance trade-off framework and illustrate this through simulations with different levels of population structure. We conclude that, unless population structure is relatively strong (*F_st_* > 0.01), the unified estimator provides the greatest power and has negligible bias due to population stratification. When population structure is relatively strong or analyses are sensitive to even tiny amounts of confounding due to population stratification, the robust estimator should be preferred.

## 2 Results

### 2.1 Including singletons in family-based GWAS

The imputation method developed in Young et al. and implemented in *snipar* only imputes missing parental genotypes for genotyped samples with at least one genotyped first-degree relative. We propose to extend the imputation to individuals without any genotyped first-degree (or any degree) relatives so that they can be included in the family-based GWAS regression. For an individual without any genotyped relatives, we have observed the two out of four of the individual’s parents’ alleles, the same as the case of a sibling pair in IBD2. Under random-mating, the imputed parental genotypes for a singleton are therefore:

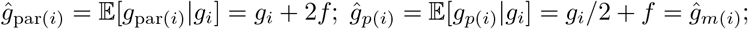

where the two unobserved alleles are imputed using the population allele frequency, *f*. The theoretical results in Young et al.[8] imply that, if the imputed parental genotypes are unbiased estimates of the true parental genotypes, then the direct effect estimates obtained when including singletons in the family-based GWAS regressions of Models 1 and 2 will be unbiased and consistent, provided that the resulting regression design matrix is not collinear. As the imputation from a singleton is linear, singleton data alone cannot be used to identify direct effects. Therefore, genotype-phenotype data from individuals with genotyped first-degree relatives, where a non-linear imputation of parental genotype(s) is possible[8], is needed in addition to singleton data to enable identification of direct effects.

Consider that we have a genotyped and phenotyped sample partitioned into two disjoint subsets: a subset with at least one genotyped first-degree relative (which we call the ‘ related sample’), where missing parental genotypes have been imputed as in Young et al.[8]; and a subset without any genotyped relatives (which we call the ‘singleton sample’), and where parental genotypes have been linearly imputed as above. We show that performing family-based GWAS on the combined sample (Figure 1), with parental genotypes replaced by their imputed values when unobserved, is equivalent to performing a multivariate, fixed-effects meta-analysis of estimates from family-based GWAS on the related sample and from standard GWAS on singletons (see Section 8.5). This equivalence enables us to easily derive theoretical results on the gain in effective sample size for direct effects from including singletons in family-based GWAS.

**Figure 1:**
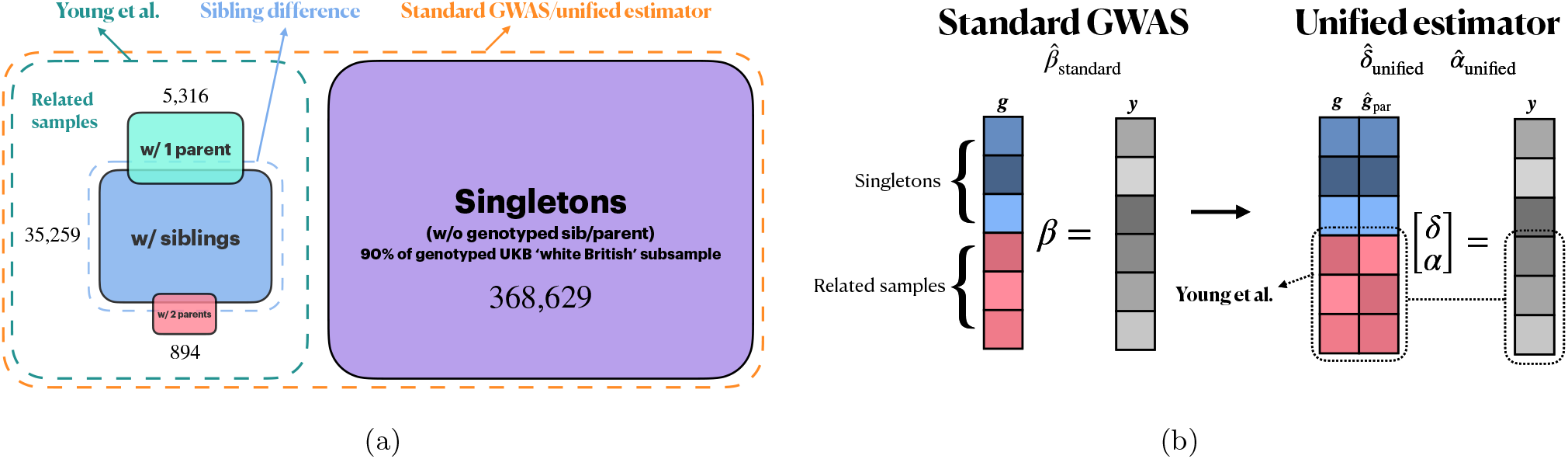
Illustrations of standard GWAS, the Young et al. estimator, and the unified estimator. *Note*. (a) We illustrate the different sample subsets used by different family-based GWAS methods and standard GWAS. We give the numbers for each subset for the UK Biobank ‘white British’ sample for illustration. The sibling difference e stimator u ses s amples w ith o ne o r m ore siblings’ genotypes observed (35,259), whereas the Young et al. estimator uses all related samples, which also include individuals with both parents’ genotypes observed (894), and those with one parent’s genotype observed (5,316); in addition to the related samples, the standard GWAS and unified estimators also use singletons (368,629). (b) Illustration of regressions p erformed by standard GWAS and the ‘ unified es timator’. Through linear imputation of parental genotypes, the unified e stimator i ncorporates s ingletons i nto t he f amily-based G WAS r egression, e nabling u se of the same sample as standard GWAS to estimate the parameter vector [*δ, α*]^*T*^. Although the design matrix for the singleton subset (in blue) in family-based GWAS is collinear, the design matrix the related sample subset (in red) is not, so the stacked design matrix is valid.

Consider the case where we have *n*_0_ independent sibling pairs whose parental genotypes are imputed, as in Young et al.[8], using phased data and IBD information, and we add to this *n*_1_ singletons with their parental genotypes linearly imputed as above. Assuming that the the correlation of the siblings’ residuals is 0 (*r* = Corr(*ϵ*_*i*1_, *ϵ*_*i*2_) = 0), including *n*_1_ singletons with their parental genotypes linearly imputed can gives an effective sample size for direct effects 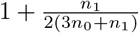 times higher than using only the *n*_0_ sibling pairs (Section 8.5.1). The theoretical gain in effective sample size converges to 50% as *n*_1_/*n*_0_ → ∞. Under these assumptions in a dataset comparable to the UK Biobank data, this implies a gain in effective sample size of ~40%.

When the correlation between siblings’ residuals is non-zero, closed-form expressions for the theoretical gain in effective sample size from adding singletons become cumbersome, but the theoretical gain can be computed numerically for particular values of *n*_0_ and *n*_1_ (see Section 8.5). Figure 2a shows the theoretical gain from adding different numbers of singletons (*n*_1_) to *n*_0_ = 20, 000 sibling pairs as a function of the correlation of the siblings’ residuals. As the correlation increases to 1, the effective sample size gain approaches 0. This is because when siblings’ residuals are perfectly correlated, noiseless estimates of direct effects can be obtained from the sibling sample alone[8].

**Figure 2:**
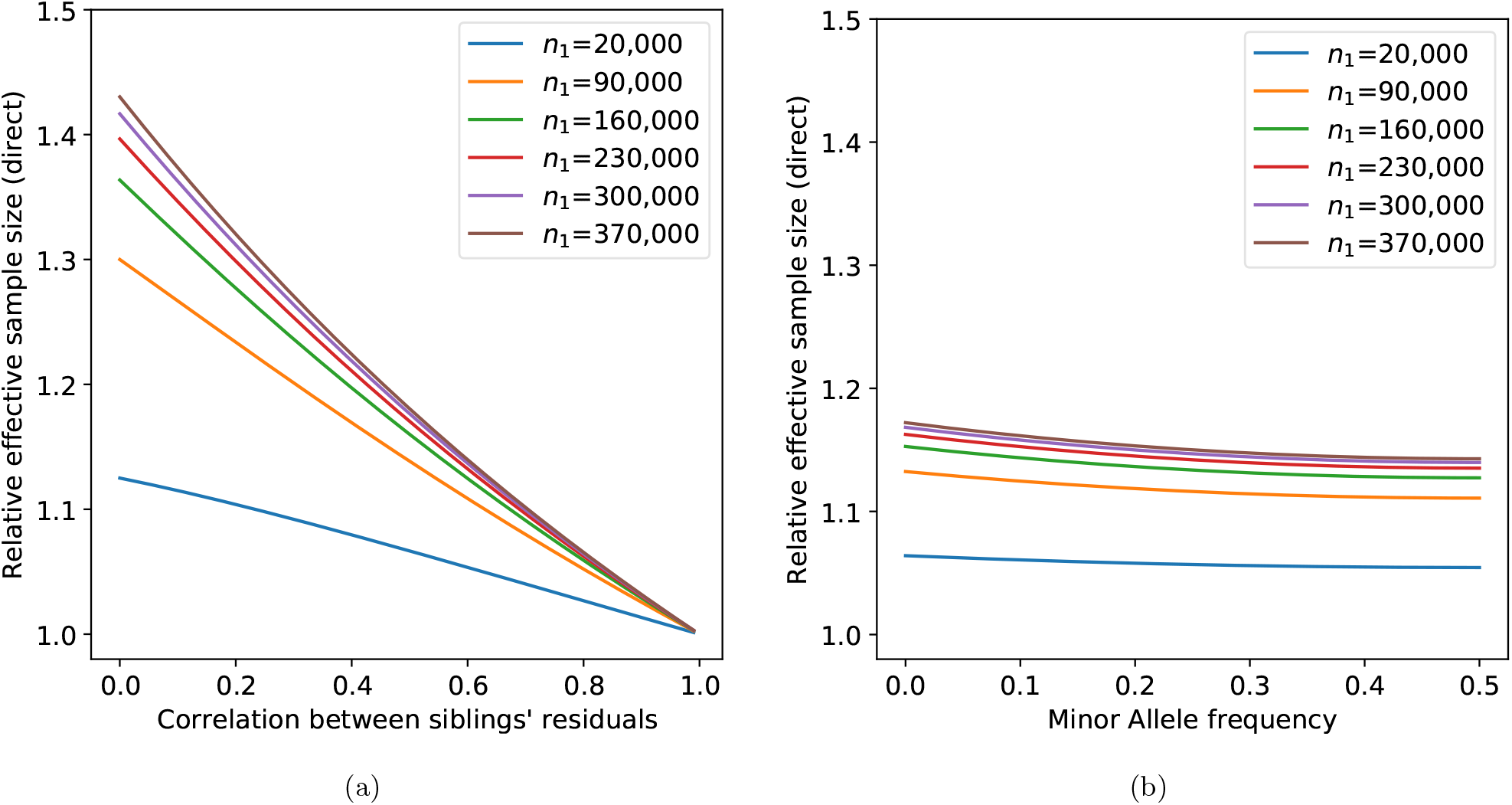
Theoretical relative effective sample sizes for 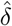. *Note*. The number of sibling pairs *n*_0_ is fixed at 20,000, which is close to the number observed in the UK Biobank ‘white British’ subsample. (a) theoretical relative effective sample sizes when parental genotype imputation for the related sample is done using phased IBD data; (b) theoretical relative effective sample sizes when parental genotypes of the related sample are imputed using unphased IBD data, in which case theoretical gain is dependent on allele frequency; sibling phenotypic correlation is fixed at 0.3.

In Section 8.5.2, we derive equivalent results for adding *n*_1_ singletons to *n*_0_ genotyped samples with one parent genotyped, where the missing parent’s genotype has been imputed using phased data as in Young et al.[8]. We find that the gain in effective sample size for direct effects is approximately 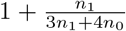 times higher than using the genotyped parent-offspring pairs alone. This implies the gain in effective sample size converges to 1/3rd as *n*_1_/*n*_0_ → ∞.

One is able to obtain an estimate of the standard GWAS ‘population effect’, *β* (which equals *δ* + *α* under random-mating), from family-based GWAS by taking the sum of the estimates of the *δ* and *α*. By performing the analysis using all genotyped samples that would normally be used in a standard GWAS (Figure 1) (i.e., restricted to have relatively homogeneous genetic ancestry), and using the same set of covariates, one can obtain an estimate of *β* similar to what one would have obtained by performing standard GWAS. We therefore call this approach to family-based GWAS, including singletons via linear imputation, the ‘unified estimator’, since it can unify family-based and standard GWAS in one analysis.

#### 2.1.1 Analysis of simulations based on UK Biobank ‘White British’ sample

We examine the proposed unified estimator on a number of populations derived from the UK Biobank ‘white British’ sub-sample. The datasets are simulated under different assumptions, including: random mating, assortative mating, population stratification, etc. Phenotypes are generated from 1,500 SNPs with randomly drawn direct and indirect effects under different correlation assumptions. The exact procedure can be found in [8]. The final output is 49,991 sibling pairs without observed parental genotypes.

To investigate the performance of the unified estimator, we randomly remove one sibling’s genotype from each sibling pair for half of the pairs, leaving us with *n*_0_ = 24, 996 genotyped sibling pairs and *n*_1_ = 24, 995 genotyped singletons. We then impute the parental genotypes of the sibling pairs with unphased IBD data using the methods described in Young et al.[8], and linearly impute those of singletons using (2.1). The results are given in Table 1. It can be seen that the observed relative effective sample size is negatively correlated with sibling correlation, with the 5th scenario observing the lowest sample size gain and the highest sibling correlation. This is consistent with the trends in Figure 2a.

**Table 1:** Results from simulated datasets. *Note*. Results for simulated phenotypes based on UK Biobank ‘White British’ sample. We imputed parental genotypes from un-phased data and IBD information, as in Young et al.[8], for *n*_0_ = 24, 996 independent sibling pairs, and we linearly imputed parental genotypes for *n*_1_ = 24, 995 singletons. Scenario 1 is simulated under random mating with a heritability of 75%; scenario 2 is generated by vertical transmission taken to equilibrium [13]: the offspring’s phenotype is affected by the parent’s phenotype with a coefficient of 0.25 and a heritability in the first generation of 50%; scenario 3 is generated by assortative mating taken to equilibrium with a phenotypic correlation between parents of *r_y_* = 0.5; scenario 4 is simulated under population stratification with 30% of the phenotypic variance explained by environmental difference between regions; in scenario 5, direct effects and IGEs from parents are perfectly correlated, *r_δ,η_*=1, and together explain 75% of the phenotypic variance. For more details please see the supplementary note in [8]. The phenotypic correlation between siblings is given in the ‘Sib. corr.’ column. The relative effective sample size when using the combined sample of sibling pairs and singletons compared to using the sibling pairs alone (with imputation) is given in the 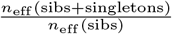 column, averaged over all SNPs with varying allele frequencies. The coefficient from regression of direct effect estimates (using the combined sample of sibling pairs and singletons) is given by the ’Coefficient 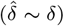’ column, and the its standard error is given in the ‘S.E.’ column.

| No. | Scenario parameters | Sib. corr. | $\frac{n_{\text{eff}}(\text{sibs}+\text{singletons})}{n_{\text{eff}}(\text{sibs})}$ | Coefficient ($\hat{\delta} \sim \delta$) | S.E. |
| --- | --- | --- | --- | --- | --- |
| 1 | $h^2=0.75$ | 0.389 | 1.074 | 0.995 | 0.009 |
| 2 | $h^2=0.5$; VT=0.25 | 0.512 | 1.060 | 0.997 | 0.012 |
| 3 | $h^2=0.5$; AM with $r_y=0.5$ | 0.386 | 1.077 | 0.993 | 0.011 |
| 4 | $h^2=0.5$; pop. strat.=0.3 | 0.549 | 1.052 | 0.999 | 0.009 |
| 5 | $h^2=0.75$; $r_{\delta,\eta}=1$ | 0.674 | 1.038 | 1.028 | 0.015 |

We compared the estimated direct SNP effects to the true causal effects. We regress the estimates onto the true values, inverse-weighted by the estimation variance. The regression coefficients and standard errors are given in the last two columns of Table 1. For all simulation scenarios, the regression coefficient is close to 1, and within two standard errors of 1, indicating little to no inflation or deflation of direct effect estimates under these scenarios.

### 2.2 Effect of population structure

We have derived the above results assuming random-mating. Here, we examine the effect of population structure on imputation and different family-based and standard GWAS estimators. As in [8], we consider an island model of population structure: the population is divided into *K* subpopulations, with random-mating within each subpopulation and no migration between them. For one locus, we denote the *k*^th^ subpopulation allele frequency as *f_k_* for *k* = 1, 2,…, *K*, and the overall population allele frequency as 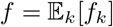, with the expectation taken over the *K* subpopulations. Parental genotype imputation that we have been considering so far relies on the overall population allele frequency. In the presence of population structure, imputation based on the overall population allele frequency *f* will be biased if *f_k_*’s are different from *f*. As a result, bias might also be introduced into the direct effect estimate. For instance, for imputation from a sibling pair with phased data, the imputed sum of parental genotypes for family *i* in subpopulation *k* is given by

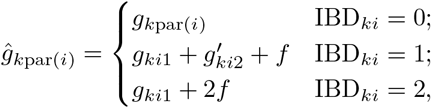

where 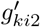 is the allele of sibling 2 that is not shared IBD with alleles inherited by sibling 1, and IBD_*ki*_ is the IBD state of the *i*th sibling pair in the *k*th subpopulation. Taking the expectation, we have 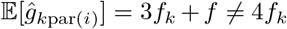. Note that we also have 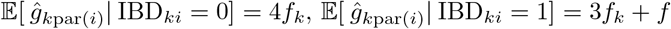, and 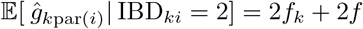. The variation in the expectation of the imputed parental genotypes across IBD states is due to the difference in number of observed parental alleles.

Consider performing family-based GWAS using siblings with parental genotypes imputed as above and with the following phenotype model:

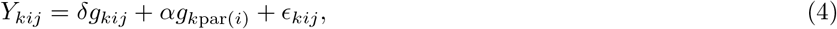

where *Y_kij_* is the phenotype of sibling *j* in family *i* from subpopulation *k*, and *ϵ_kij_* is uncorrelated with both proband and parental genotype. Young et al. [8] showed that performing the regression with *g*_*k*par(*i*)_ replaced with 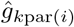 (as defined above) gives, in limit, 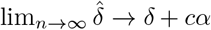, where *c* is a function of Wright’s *F_st_*, defined as *F_st_* = Var_*k*_(*f_k_*)/[*f* (1 − *f*)] in this model. When *F_st_* is small, *c* ≈ *F_st_*/2, implying the bias will be negligible for European genetic ancestry samples, where *F_st_* has been estimated to be on the order of 10^−3^[14]. In contrast, under this model of population structure, the population effect is 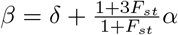.

#### 2.2.1 Population structure robust estimator

Young et al. proposed an alternative, imputation-based estimator for sibling pair data that they argued should be robust to population structure. This estimator analyzes sibling pairs in IBD0 and IBD1 separately and then meta-analyzes the direct effect estimates. The intuition here is that the bias introduced into direct effect estimates is because the bias in the imputed parental genotype varies across IBD states, creating excess variance in the imputed parental genotype that is correlated with population structure. Therefore, by analysing sibling pairs separately depending upon IBD state, the excess variance in the imputed parental genotype due to population structure is removed. However, for siblings in IBD2 (as for singletons), the imputed parental genotype is collinear with the siblings’ genotypes, implying that direct effects cannot be identified based upon siblings in IBD2 alone. Sibling pairs in IBD2 are thus ignored in this estimator. Since they are genetically identical, sibling pairs in IBD2 also do not contribute to the sibling difference estimator.

Young et al. proved that fitting Model 1 for sibling pairs in IBD0 or IBD1 (with parental genotypes imputed as above) gives consistent estimates of direct effects in this model of population structure[8]. When sibling pairs are in IBD0, all four parental alleles are observed, so this situation is similar to having both parents genotyped, except the parental origin of alleles is (in general) unknown. For siblings in IBD1, one allele in each sibling is an allele that was not transmitted to the other sibling, and these non-transmitted alleles provide the information required to obtain consistent estimates of direct effects[8]. By estimating direct genetic effects separately for sibling pairs in IBD0 and IBD1, and then meta-analyzing the result, one can obtain a consistent estimator of direct effects in a structured population. Young et al.[8] showed that this estimator is more powerful than the sibling difference estimator, having a relative effective sample size is 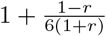 times greater, where *r* is the correlation of siblings’ residuals. However, the robustness of this estimator comes at the cost of a smaller effective sample size for estimation of direct effects than the primary estimator considered in Young et al.[8], which includes sibling pairs in IBD2.

Here, we generalize the robust estimator proposed by Young et al. so that it can handle all possible data types. We generalize this by partitioning the sample not on IBD state but on which parental alleles that were not transmitted to the focal, phenotyped individual (proband) we have observed. We give the partition of the sample in Table 2, and we prove that the regressions in these subsamples give consistent estimates of direct effects (Section 8.6). Thus, we propose a generalized robust estimator that is able to yield consistent estimates of direct effects in structured populations by performing separate analyses on the four groups and performing a fixed-effects, inverse-variance weighted meta-analysis of the direct effect estimates. The sibling pair based estimator proposed in Young et al.[8] can be seen as a special case, because sibling pairs in IBD0 and IBD1 belong to the ‘both NT’ and ‘one NT’ groups respectively (Table 2). This further increases the power advantage of the robust estimator over the sibling difference estimator (which is less powerful even for sibling pair data) by enabling inclusion of additional samples, such as samples with genotyped parent(s) but without genotyped siblings.

**Table 2:** Partition of sample with at least one non-transmitted allele observed. *Note*. For the robust estimator, we partition the sample with at least one non-transmitted (NT) parental allele observed into four groups. By performing the regressions separately in each group, we obtain consistent estimates of direct effects from each group, even when there is population structure so that imputation based on the overall allele frequency is biased (Section 8.6). For the regression column, *y_ij_* is the phenotype of sibling *j* in family *i*; *g_ij_* the corresponding genotype; *g*_*p*(*i*)_ and *g*_*m*(*i*)_ are the paternal and maternal genotypes; *g*_par(*i*)_ = *g*_*p*(*i*)_ + *g*_*m*(*i*)_. A caret indicates a genotype that has been imputed from phased data as in Young et al.[8]; e.g., 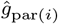 refers to the imputed sum of parental genotypes.

| Group | Example data types | NT alleles observed | Regression |
| --- | --- | --- | --- |
| Maternal NT | mother-child pairs<br>mother and a sibling pair in IBD2 | maternal | $y_{ij} \sim g_{ij} + \hat{g}_{p(i)} + g_{m(i)}$ |
| Paternal NT | father-child pairs<br>father and a sibling pair in IBD2 | paternal | $y_{ij} \sim g_{ij} + g_{p(i)} + \hat{g}_{m(i)}$ |
| Both NT | sibling pairs in IBD0<br>genotyped trios | paternal and maternal | $y_{ij} \sim g_{ij} + g_{\text{par}(i)}$ |
| One NT | sibling pairs in IBD1 | paternal or maternal | $y_{ij} \sim g_{ij} + \hat{g}_{\text{par}(i)}$ |

#### 2.2.2 Comparison of estimators in structured populations

Here, we examine the power (measured by effective sample size) and bias of the different estimators (unified, robust, Young et al., sibling difference, and standard GWAS; see Table 3 for summary) in simulations with different levels of population structure, as measured by Wright’s *F*_ST_. We simulated populations of sibling pairs, where the population was divided into two equally sized subpopulations, and the allele frequencies for the two subpopulations for 20,000 SNPs were simulated from the Balding-Nichols model[15] (Section 8.3). For simulation results including the unified estimator, we simulated 2,000 sibling pairs and 18,000 singletons in each subpopulation. For simulations without the unified estimator, we simulated 20,000 sibling pairs in each subpopulation. The phenotypes were generated such that mean difference between the two subpopulations explained 50% of the overall phenotype variance, and the remaining 50% of the phenotype variance was generated by random Gaussian noise, implying a correlation between siblings’ phenotypes of 0.5. There are no causal effects (or direct effects) of the genotypes in this simulation, so any deviation of the effect estimates from what would be expected under the null distribution is evidence of bias due to population stratification.

**Table 3:** Summary of estimators.

| Estimator | Data types used | Procedure |
| --- | --- | --- |
| Sibling difference | phenotyped and genotyped samples with at least one genotyped sibling | regress sibling phenotypic differences onto genetic differences, or regression of phenotype onto deviation of genotype from mean among siblings[10] |
| Robust | phenotyped and genotyped samples with at least one observed non-transmitted parental allele (see Table 2) | perform separate family-based GWAS regressions on different data types and meta-analyze the direct effect estimates |
| Young et al. | phenotyped and genotyped individuals with at least one genotyped first-degree relative | family-based GWAS (Models 2 and 1) with observed and/or imputed parental genotypes |
| Unified | all phenotyped and genotyped samples (with or without genotyped relatives) | perform Mendelian imputation for related samples and linear imputation for singletons, and conduct family-based GWAS (Models 2 and 1) on the combined sample |
| Standard GWAS | all phenotyped and genotyped samples (with or without genotyped relatives) | regress proband phenotypes onto proband genotypes |

We compute the mean of the squared of Z-scores, i.e., 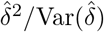 (or 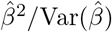) for standard univariate GWAS), of the 20,000 estimated SNP effects produced by different estimators, which should be 1 in expectation under the null, and will be above 1 in expectation if there is bias due to population stratification. While a mean *Z*^2^ statistic greater than 1 is a common measure of inflation in GWAS[16, 17], this statistic is not a completely fair way to compare the biases due to stratification across estimators that have different sampling variances: for example, for estimators with the same bias but different sampling variances, the estimator with the smaller sampling variance would be expected to produce larger *Z*^2^ statistics on average. Hence, we also look at the non-sampling variance of an estimator *ζ*, 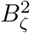, across the *L* = 20, 000 SNPs, which we estimate as

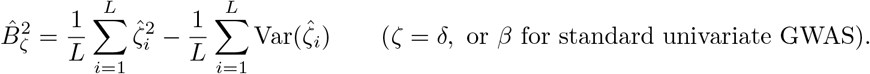

Consider that an estimator for a SNP *i*, 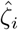, has expectation *b_ζi_* due to bias from population stratification, then the expectation of the non-sampling variance estimator is

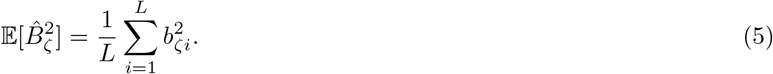

Thus, 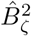 gives an estimate of the magnitude of bias due to population stratification that can be fairly compared across estimators. We give values for 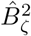 relative to the maximum observed, which is the non-sampling variance of the standard GWAS estimator when *F*_ST_ = 0.1.

We display the results for populations with *F_st_* = 0, 10^−3^, 10^−2^, 10^−1^ in Figure 3, where *F_st_* = 10^−3^ is roughly the level of differentiation between neighbouring European populations[14], and *F_st_* = 10^−1^ is roughly the level of population differentiation between European ancestry and East Asian ancestry populations[18]. As expected from theory, the sibling difference and the robust estimators have no detectable bias from population stratification for any level of *F_st_*, and the standard GWAS estimator has the most bias, with statistically significant bias for *F_st_* ≥ 10^−3^. The unified and Young et al. estimators do not have detectable bias except for *F_st_* = 0.1, with the unified estimator having greater bias than the Young et al. estimator. This is expected since the unified estimator includes a large sample of singletons, where the two unobserved parental alleles are imputed using the overall population frequency, which is biased in a structured population.

**Figure 3:**
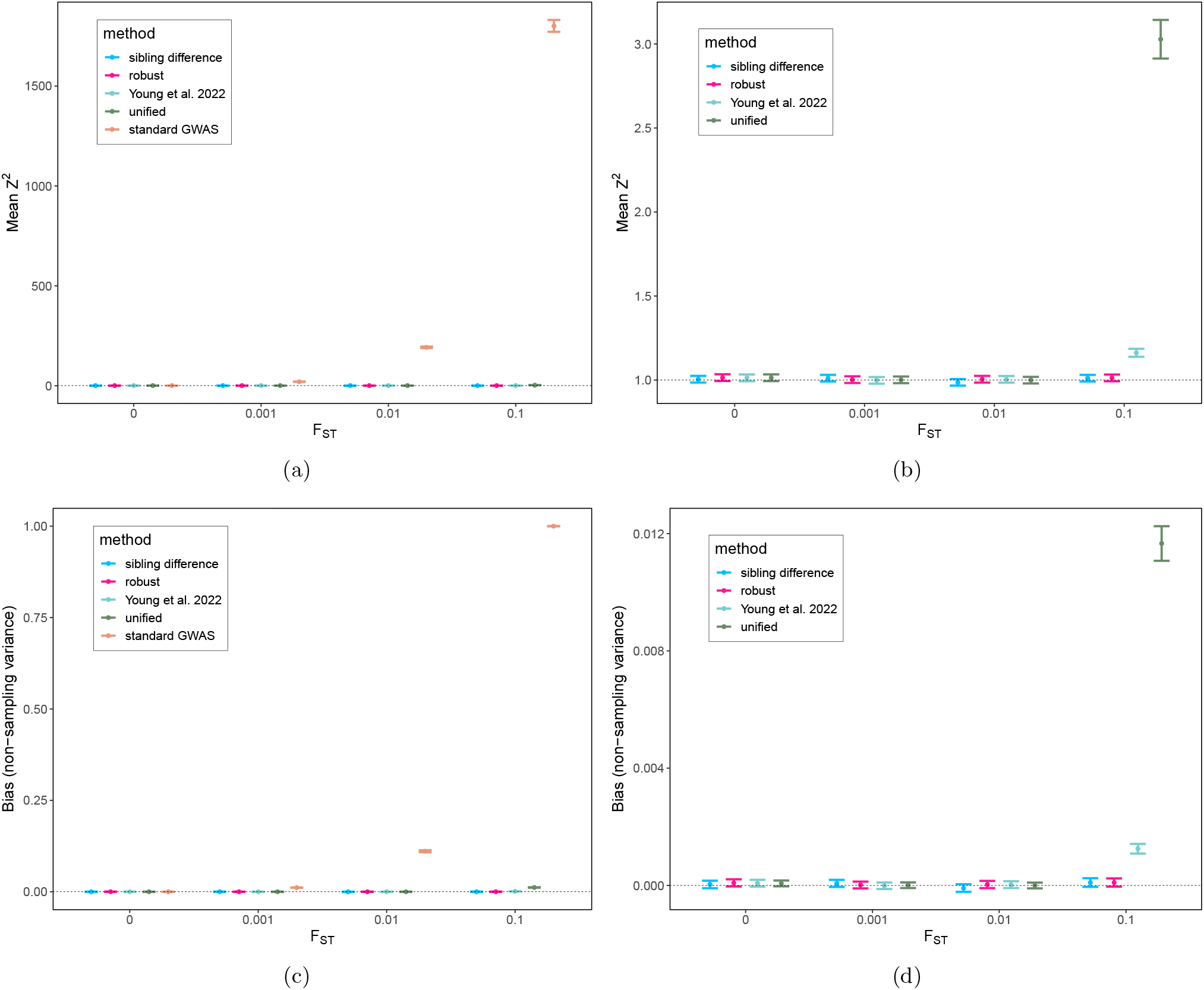
Comparison of bias due to population stratification for different estimators and different levels of population structure. *Note*. Error bars display 95% confidence intervals. (a) Mean of squared Z-statistics across 20,000 SNPs for the four estimators, which are expected to be above 1 (dashed-line) when there is bias due to population stratification; (b) same as (a) but with the standard GWAS removed; (c) non-sampling variances (see 2.2.2) of the estimators relative to the ‘maximum’ observed (for standard GWAS with *F_st_* = 0.1), which gives a measure of the magnitude of bias due to population stratification, with values above 0 indicating bias; (d) the same as (c) but with the standard GWAS removed.

#### 2.2.3 Bias-variance trade-off of estimators

We compare the estimators (Table 3) in a bias-variance trade-off framework. We display results for a scenario mimicking the UK Biobank in Figure 4: around 10% of the sample has a genotyped sibling, and the remaining 90% are singletons. We use this to illustrate the gain in power from including the singletons in the ‘unified estimator’. We display results for a sibling pair only scenario in Figure 5. See Section 8.3 for simulation details. We calculate the effective sample size relative to that of the sibling difference estimator. Thus, an effective sample size greater than 1 means higher precision (and statistical power) than the sibling difference method. Bias is evaluated as the non-sampling variance relative to the ‘maximum’ observed, as defined in the previous section.

**Figure 4:**
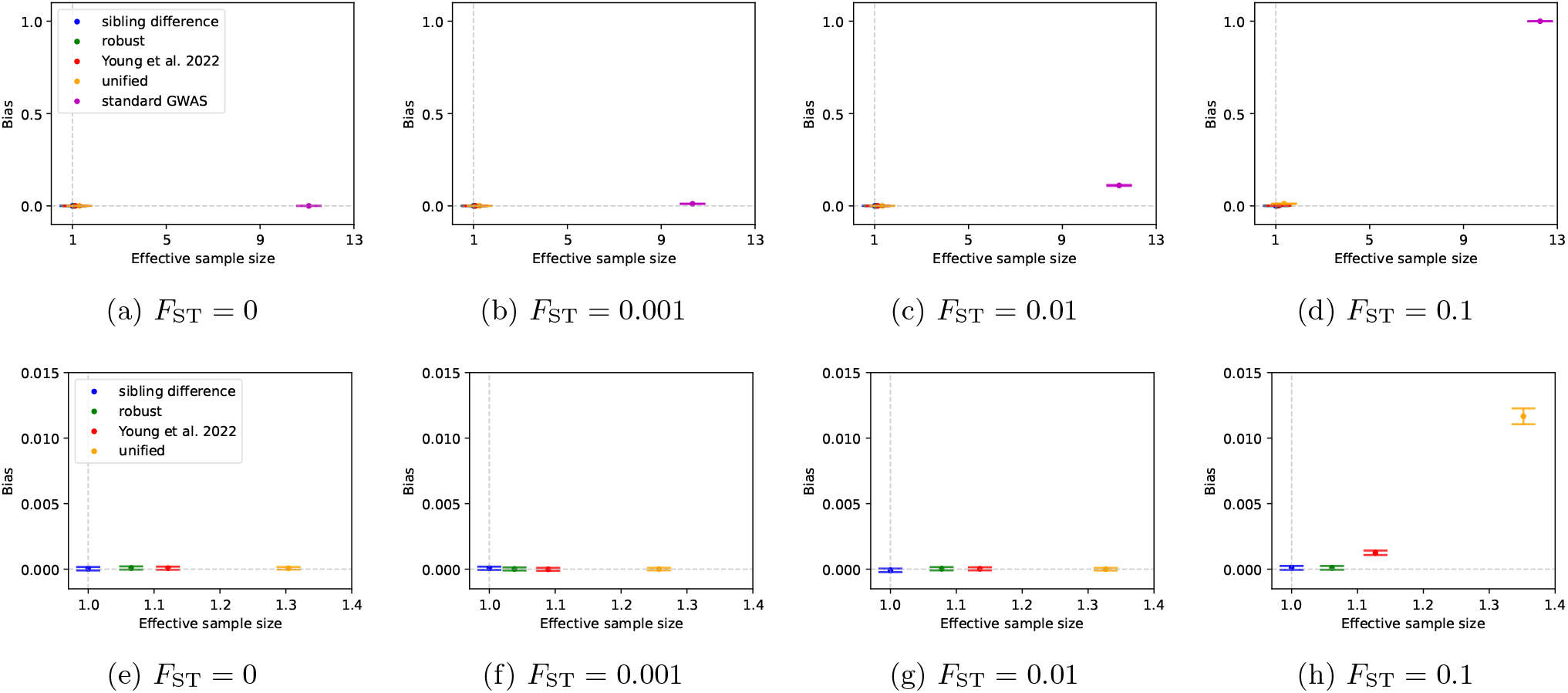
Bias-variance tradeoff on simulated sibling pairs and singletons. *Note*. The simulated datasets in Figure 3 are used for this demonstration: 2,000 independent sibling pairs and 18,000 singletons in each of two subpopulations differentiated at the level given by *F_st_*. Effective sample size is defined relative to that of the sibling difference estimator. Bias is measured as the non-sampling variance (2.2.2) relative to the maximum observed, which is for standard GWAS with *F_st_* = 0.1. (a)-(d) bias-variance tradeoff comparison for the sibling difference method, robust estimator, Young et al., unified estimator, and standard GWAS; (e)-(h) the same as (a)-(d) but with the population effect removed for scale.

**Figure 5:**
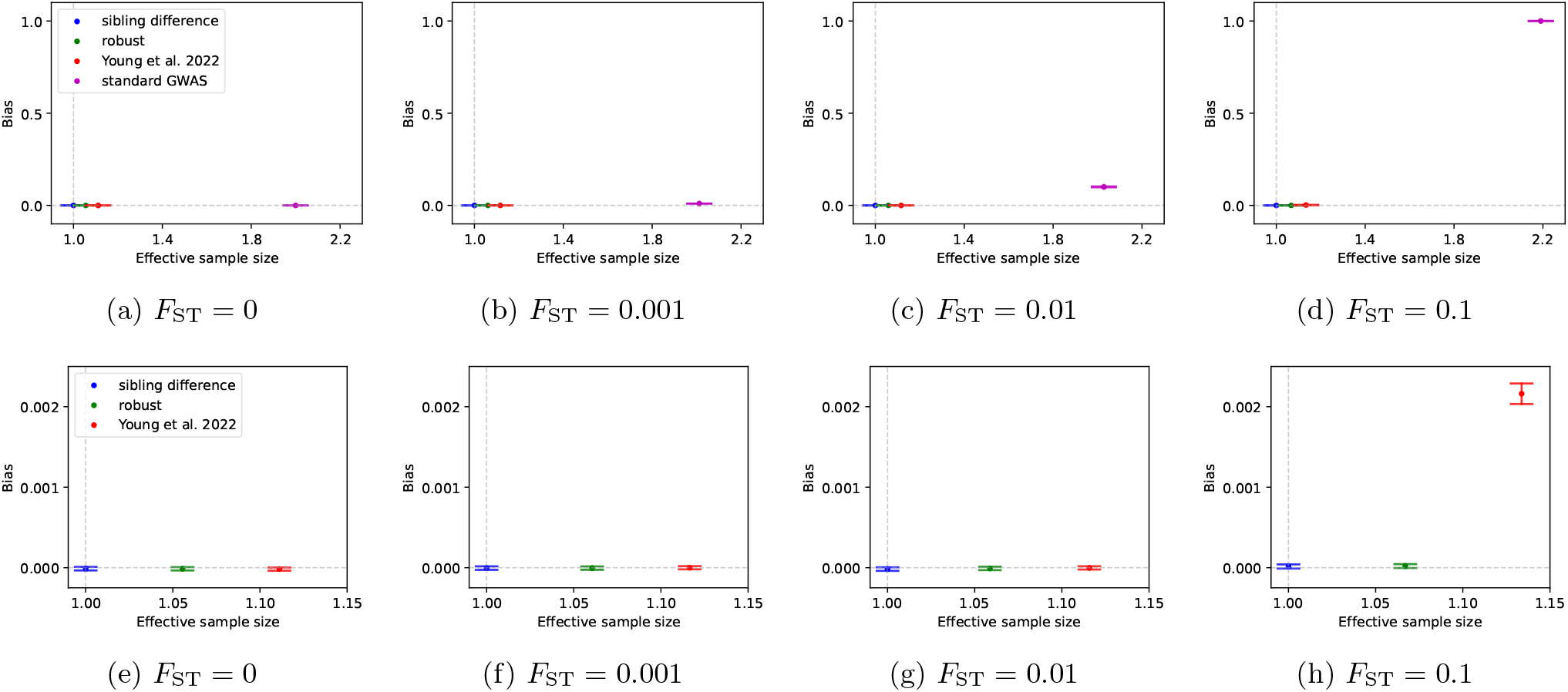
Bias-variance tradeoff on simulated sibling pairs. *Note*. We simulated populations and phenotypes as in Figure 4, except that we simulated 40,000 sibling pairs (20,000 in each subpopulation). In this case, the unified estimator and the Young et al. estimator are equivalent, because there are no singletons. Effective sample size is defined relative to that of the sibling difference estimator. Bias is defined as the non-sampling variance (2.2.2) relative to the ‘maximum’, the non-sampling variance of the standard GWAS estimate when *F*_ST_ = 0.1. (a)-(d) bias-variance tradeoff comparison for the sibling difference method, robust estimator, Young et al. estimator, and standard GWAS; (e)-(h) the same as (a)-(d) but with the population effect removed for scale.

The simulations confirm theoretical expectations. The sibling difference estimator is robust to population structure, but it is less powerful than the imputation based ‘robust estimator’, which gains power by incorporating information on non-transmitted parental alleles deduced by Mendelian imputation for sibling pairs in IBD0 and IBD1. By also considering sibling pairs in IBD2, the Young et al. estimator can further boost statistical power, but introduces a slight bias due to population structure that becomes detectable when *F*_ST_ > 0.01. The ‘unified estimator’ incorporates information from singletons in addition to the samples used by the Young et al. estimator, gaining power; however, this power comes at the cost of increased bias due to population structure, although this only becomes apparent when *F*_ST_ > 0.01. The standard GWAS estimator has much greater effective sample size than the other methods, but this comes at the cost of much greater bias in structured populations.

The simulation results show that, out of the family-based estimators considered, the unified estimator has the greatest power and has negligible bias due to population structure in relatively homogeneous samples (e.g., *F*_ST_ ≤ 0.01), such as the European genetic ancestry samples often used in GWAS. However, in samples with relatively strong structure (*F*_ST_ ≥ 0.01), the ‘robust estimator’ should be preferred. The greater effective sample size of the ‘robust estimator’ over the sibling difference estimator is calculated here for a sample of sibling pairs only, and where siblings have a phenotypic correlation of 0.5. The advantage of the ‘robust estimator’ will be larger in samples including genotyped parents or families with more than two genotyped siblings, and will be larger when siblings’ phenotypes (more precisely, their residuals) have correlation less than 0.5.

### 2.3 Linear mixed model inference

The above theory and simulation results have assumed we have independent sibling pairs and independent singletons, i.e. the only relatedness in the sample is between siblings. For the simulation results, we used the same LMM as in Young et al., which models the mean difference in phenotype between sibships as a random effect, thereby accounting for residual correlations between siblings’ phenotypes and ensuring estimates of direct effects are statistically efficient[8]. Here, we develop a LMM that generalizes the LMM used in Young et al. and the LMM implemented in *fastGWA*[12], which is specified by a sparse genetic relatedness matrix (GRM). This approach ensures that residual correlations between siblings are modelled properly, ensuring statistically efficient estimates of direct effects are obtained, while also modelling residual correlations between all pairs related above some threshold, thereby ensuring statistically efficient estimates with accurate standard errors errors are obtained when more complex relatedness is present in the sample[12].

Stacking all observation vertically, for a dataset with *N* individuals in *n* families, the model is

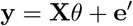

where **y** is the *N* × 1 phenotype vector; **X** is the *N* × *c* matrix specifying the fixed effects, where the columns of **X** depend upon the covariates and estimator being used; *θ* is the corresponding vector of fixed effects; and **e′** is a random vector, which we specify below. For example, if we are fitting Model 2 without additional covariates, *θ* = [*δ, α_p_, α_m_*,] ^⊤^, and **X** has columns giving proband, (imputed or observed) paternal, and (imputed or observed) maternal genotypes. The random vector **e′** is specified as

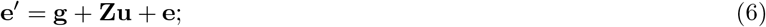

where

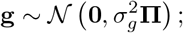

where **Π** is the (sparse) genetic relatedness matrix, and 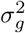 is the corresponding variance parameter; **Z** is a *N × n* sibship-indicator matrix, the *kl*th entry is 1 if the *k*th individual is in sibship *l* and 0 otherwise, and **u** is an *n*×1 normally distributed sibship-specific mean vector:

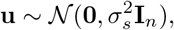

where 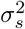 is the sibship covariance parameter. The the sibship covariance component **s** is thus also normally distributed:

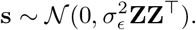

The residual variance vector has distribution:

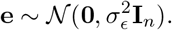

Therefore, the variance-covariance matrix of **y**|**X** is given by 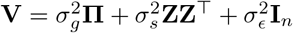.

The relatedness matrix, **Π**, can be either a SNP-based genomic relationship matrix (GRM) or a relatedness matrix computed from IBD segments, such as output by KING[19]. By setting elements of **Π** below a certain threshold, usually 0.05, to zero, the sparsity of the **V** matrix can be exploited so that REML inference of variance components, and generalized least-squares inference of *θ* given the REML variance components, is computationally feasible even for large-scale biobanks[12](see Section 8.1).

### 2.4 The unified estimator increases power for estimating direct effects in the UK Biobank

To demonstrate the increase in effective sample size for direct effects from the unified estimator, we apply the unified estimator and the Young et al. estimator to the UK Biobank ‘white British’ subsample. We used the LMM outlined above, with the sparse genetic relatedness matrix derived from IBD segments inferred by KING, with the relatedness threshold set at 0.05.

We analyzed 10,911 SNPs on chromosome 22 for 20 health and behavioral phenotypes (Table 4; see Section 8.4 for analysis steps). We compare the effective sample size for direct effects from the unified estimator relative to the Young et al. estimator as a function of phenotypic correlation between siblings. Figure 6 and Table 4 show gains in effective sample size between 25.9% and 42.7%, depending on the phenotype, with phenotypes with higher correlations between siblings exhibiting less gain, as expected from theory (Figure 2a).

**Table 4:** Results from analysis of UK Biobank data. *Note*. We list phenotypic correlations of siblings (‘sib. corr.’) and effective sample sizes of the unified estimator relative to the Young et al. estimator (see Table 3 and Figure 6). Abbreviations: HDL, high density lipoprotein; BMI, body mass index.

| Phenotype | Sib. corr. | Relative effective sample size |
| --- | --- | --- |
| Blood glucose | 0.0792 | 1.4144 |
| Subjective well-being | 0.0921 | 1.4235 |
| Number of children (male) | 0.1297 | 1.4266 |
| Neuroticism | 0.1376 | 1.3945 |
| Self-rated health | 0.1492 | 1.3769 |
| Number of children (female) | 0.1505 | 1.4017 |
| Diastolic blood pressure | 0.1580 | 1.3721 |
| Systolic blood pressure | 0.1587 | 1.3733 |
| Drinks-per-week | 0.1692 | 1.4162 |
| Non-HDL cholesterol | 0.1764 | 1.3702 |
| Cigarettes-per-day | 0.1830 | 1.4130 |
| Ever-smoked | 0.1838 | 1.3567 |
| Myopia | 0.1960 | 1.3567 |
| Household income | 0.2411 | 1.3504 |
| BMI | 0.2782 | 1.3238 |
| HDL cholesterol | 0.2807 | 1.3356 |
| Age-at-first-birth (women) | 0.2989 | 1.3813 |
| Cognitive ability | 0.3093 | 1.3606 |
| Educational attainment (years) | 0.4023 | 1.2886 |
| Height | 0.4978 | 1.2592 |

**Figure 6:**
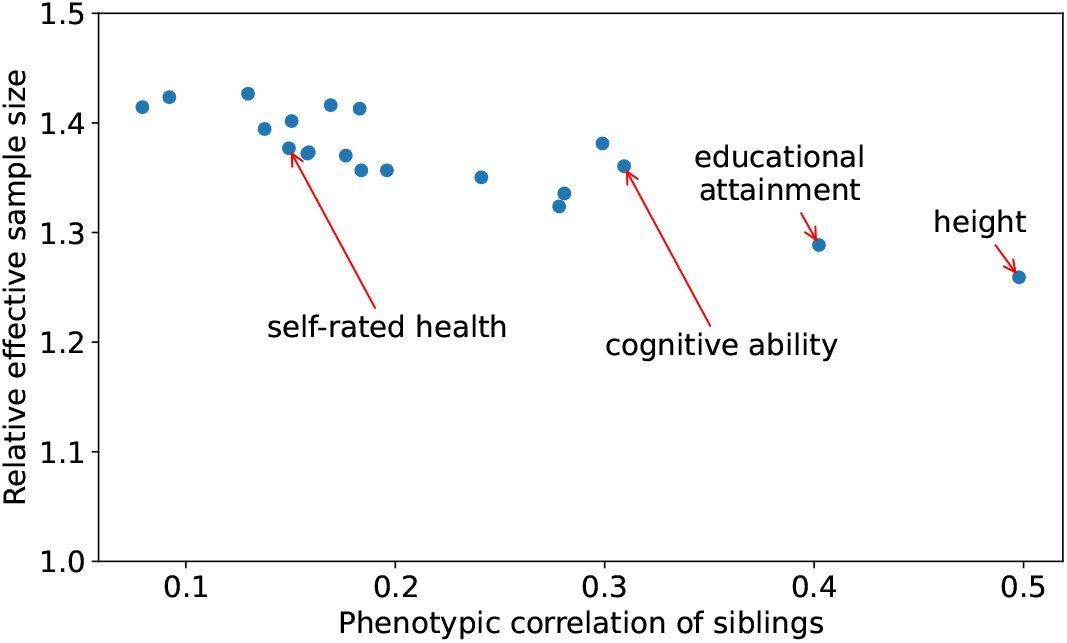
Effective sample size gain from adding singletons to family-based GWAS in UK Biobank *Note*. For each phenotype, we look at the median effective sample size of the unified estimator relative to the Young et al. estimator (for direct effects) across 10,911 SNPs on chromosome 22.

## 3 Discussion

Most genome-wide association studies (GWAS) published to date have been conducted in samples with relatively homogeneous ancestry without using genotypes of siblings or parents to remove confounding, a method we call ‘standard GWAS’. It is well established that standard GWAS often fails to eliminate all confounding, e.g., due to gene-environment correlation (including indirect genetic effects and population stratification) and assortative mating. The consequences of this bias include: 1) overestimation of heritability and the traits’ shared genetic architectures[20–22]; 2) spurious inferences of disease causes by Mendelian Randomization[23]; 3) bias in polygenic indexes (PGIs, also called polygenic scores) that contributes to the drop in predictive accuracy when predicting across genetic ancestries[24–26]; and 4) spurious inferences of natural selection[6, 10, 27]. The remedy for these problems is more precise estimates of direct effects of genome-wide SNPs from family-based designs, which remove confounding, across diverse genetic ancestries. While collecting more genotype data on families is essential for this, ensuring we have methods that make the most of available data is of crucial importance.

Here we build upon the family-based GWAS framework introduced by Young et al.[8], which treats parental genotypes as missing data and imputes them based on Mendelian Laws, substituting the imputed parental genotypes for the true parental genotypes in Models 2 and 1. We introduced novel family-based GWAS estimators that take a step towards maximizing the scientific returns from available genotype data: 1) the ‘unified estimator’ that maximizes power for estimation of direct genetic effects in relatively homogeneous samples (*F_st_* <= 0.01), such as the European genetic ancestry samples typically used to date in GWASs; and 2) the ‘robust estimator’ that is robust to population structure, so can be applied to genetically diverse samples (*F_st_* > 0.01) without introducing bias, and has greater power than alternative estimators that are robust to population structure. The ‘robust estimator’ may also be preferred for analyses of relatively homogeneous samples that are sensitive to even tiny amounts of bias due to population stratification.

We implement our estimators in a linear mixed model (LMM) framework that accurately models phenotypic correlations between siblings while accounting for sample relatedness beyond sibling pairs through a sparse genetic relatedness matrix (GRM). We use sparse linear algebra routines to enable the estimators to be applied to large-scale biobanks, which we demonstrate through application of the unified estimator to UK Biobank data. We showed that the unified estimator increases effective sample size for direct effects by between 25% and 40%, depending upon phenotype, compared to the method of Young et al.[8], which only uses samples with genotyped first-degree relatives. An advantage of the unified estimator is that, in addition to producing more precise direct effect estimates than alternative approaches, it also produces estimates of the standard GWAS ‘population effect’ that are competitive with state-of-the-art standard GWAS methods, such as *fastGWA*[12]. Both standard and family-based GWAS summary statistics, and their joint sampling distribution, can therefore be obtained from one analysis with the unified estimator.

We compared the estimators (Table 3) in a bias-variance trade-off framework for simulated populations exhibiting different levels of population structure. From this, we can order the different estimators based on increasing effective sample size (or statistical power): sibling difference, robust, Young et al., unified, and standard GWAS. This reflects the ordering in terms of bias, except for the sibling difference and robust estimators, that are both robust to population structure, so do not exhibit any bias.

We have focused on the theoretical properties of the estimators and their performance in simulations. Future work will further examine their performance on real world data: in particular, the performance of different family-based GWAS estimators when applied to genetically diverse samples.

## 4 Data availability

Summary statistics from different estimators applied to UK Biobank data will be made available on publication. Applications for access to the UKB data can be made on the UKB website (http://www.ukbiobank.ac.uk/register-apply/).

## 5 Code availability

The code implementing family-based GWAS as in Young et al.[8] is available in the *snipar* software package: https://github.com/AlexTISYoung/snipar/

The code for preforming family-based GWAS with the ‘robust’ and ‘unified’ estimators is under development and available as a development branch of *snipar* at https://github.com/AlexTISYoung/snipar/tree/fgwas_v2_merge.

## 6 Acknowledgements

The study was supported by Open Philanthropy (010623-00001 and 2019-198171) and the National Institute on Aging/National Institutes of Health through grants R24-AG065184 and R01-AG042568 (to the University of California, Los Angeles). This research has been conducted using the UK Biobank Resource under Application Number 11425.

## 7 Author Contributions

A.I.Y. conceived the study. A.I.Y. and J.G. derived theoretical results. A.I.Y. and J.G. designed the simulations. J.G. performed the simulations. S.M.N. wrote the imputation code. J.G. analyzed the UK Biobank data. A.I.Y., J.G., and D.J.B. wrote the manuscript.

## 8 Methods

### 8.1 Variance component estimation

The variance component parameters 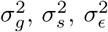 are estimated by maximizing the REML log likelihood function:

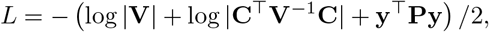

where **C** is the design matrix of fixed covariates, and

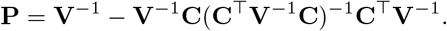

If no fixed covariate is controlled for, **C** will be a column vector of all 1’s.

If the relatedness matrix **Π** is dense, then **V** is dense, leading to resource-demanding and time-consuming computation. To reduce the computational burden, we follow Jiang et al.[12] and zero out entries in **Π** with relatedness below a default threshold of 0.05. This results in a highly sparse covariance matrix, enabling the use of more efficient sparse matrix algorithms for likelihood evaluation. By using a gradient-free optimizer, REML variance component estimation can be done in just a few minutes for datasets as large as the UK Biobank. Another possible benefit is that, by considering only close relatives, the correlations between close relatives are modelled more accurately than when using a SNP based relatedness matrix including relatedness between all pairs[12, 22].

With a sparse **V**, we compute **V**^−1^**y** and **V**^−1^**C** using a sparse LU solver in SciPy[28], without explicitly computing **V**^−1^. Then variance component parameters are optimized using the gradient-free L-BFGS algorithm [28].

One can choose to model only the sibship variance component and the residual variance component as in [8], which also results in a sparse **V** matrix, so the same computational procedure can be used in this case.

### 8.2 Estimating SNP effects

Genotype data is usually split into multiple files by chromosome, and can be processed sequentially. For each genotype file, SNPs are divided into small batches and analyzed in parallel, where the number of batches is dependent on the number of individuals and the number of cores users want to leverage.

To include covariates in the genome-wide estimaation of SNP effects, we project both genotypes and phenotypes into the space orthogonal to the space spanned by the covariates, as in BOLT-LMM[29]:

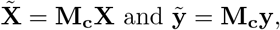

where **M_c_** = **I**_*N*_ − **C**(**C**^⊤^**C**)^−1^**C**^⊤^ is the projection matrix. Then the effect estimates are given by

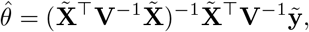

where

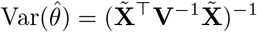

is the sampling variance-covariance. By the Frisch-Waugh-Lovell Theorem, this gives estimates of the SNP effects that are equivalent to performing the joint-regression on the covariates and the proband and (imputed) parental genotype(s).

### 8.3 Simulating structured populations

For different levels of *F*_ST_, we generated 2 subpopulations with different fixed subpopulation effects, each with 20,000 sibling pairs. We simulated 20,000 SNPs from binomial distributions, where subpopulation allele frequencies were drawn from the Balding-Nichols model[15]: 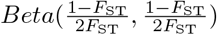. We then generated phenotypes with 50% of the phenotypic variance attributed to fixed subpopulation means, and the remaining 50% attributed to random Gaussian noise. There are therefore no nonzero direct effects in these simulations.

For the simulations involving the unified estimator, we sought to mimic the fact that large biobank datasets such as the UK Biobank consist mostly of singletons. We randomly removed one sibling from 90% of the sibling pairs to obtain 18,000 singletons and 2,000 sibling pairs. The sibling difference, robust, and Young et al. estimators were applied to the 2,000 sibling pairs, while the unified estimator and standard univariate GWAS were applied to the combined sample of 18,000 singletons and 2,000 sibling pairs.

We also examine the performance of the estimators in a sibling pair only scenario: i.e., 20,000 genotyped and phenotyped sibling pairs in each subpopulation. We applied the estimators to the resulting 40,000 sibling pairs. Note that in this scenario, there are no singletons, and the unified estimator is equivalent to the Young et al. estimator.

### 8.4 Analysis of UK Biobank data

We filtered the genotyped samples to 408,254 individuals who are of ‘white British’ genetic ancestry (as determined by UK Biobank[11]) with no excessive relatives, no sex chromosome aneuploidy, and not identified as outliers in heterozygosity and genotype missingness. We used the phased haplotypes for the UK Biobank genotyping array SNPs provided as part of the UK Biobank data release. We filtered out variants with minor allele frequency less than 0.01, with Hardy-Weinberg equilibrium exact test p-value less than 1e-6, and with imputation (INFO) quality score less than 0.99, resulting in 658,720 SNPs (10,911 SNPs in chromosome 22). We inferred IBD segments shared between siblings and performed imputation using *snipar* [8]. We then imputed the parental genotypes of the remaining singletons with (2.1). We fitted model (2) for each SNP, substituting imputed parental genotypes for observed parental genotypes when not available.

The UK Biobank phenotypes are as described in Okbay et al. 2022[26].

### 8.5 Theoretical effective sample size for the unified estimator

In section 2.1 we claim that performing family-based GWAS on the combined sample yields the same results as meta-analyzing results from family-based GWAS on the related sample and standard univariate GWAS on singletons. To see that, we first look at a general case where we are given two sets of effect estimates **z**_0_ and **z**_1_ obtained from independent datasets, with

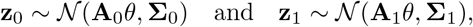

where **A**_*i*_’s are some linear transformations and **Σ**_*i*_’s are estimation covariance matrices. By [8], the MLE of the true effect sizes *θ* is given by

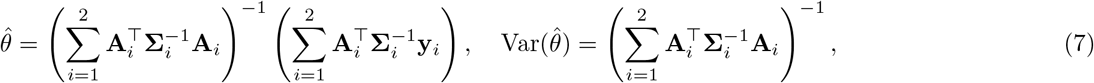

if 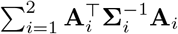 is invertible. Now suppose we have 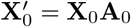 and 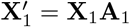, which are transformed from raw design matrices **X**_0_ and **X**_1_. Combining them into one dataset, we get

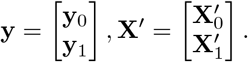

Then we have

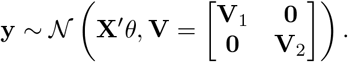

Thus,

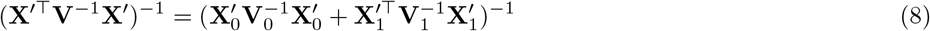

and

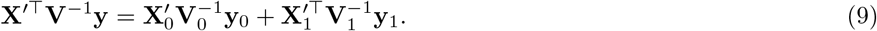

Suppose we want to estimate the direct effect and the average non-transmitted coefficient. So for the related sample, we have **A**_0_ = **I**, since we have enough information to estimate both parameters. For singletons, since the imputed parental genotype is a linear function of the proband’s genotype, the estimated effect will be the sum of the two effects, meaning that **A**_1_ = [1 1]. Also, assume that we only model the sibling variance component and the residual variance component. This gives 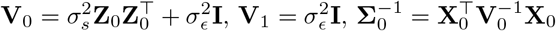 and 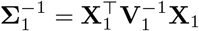, since there is no sibling pair in the sample of singletons. Combining the above, we have

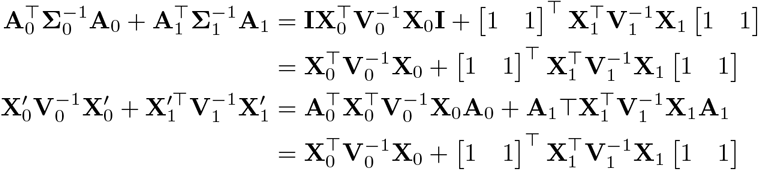

and

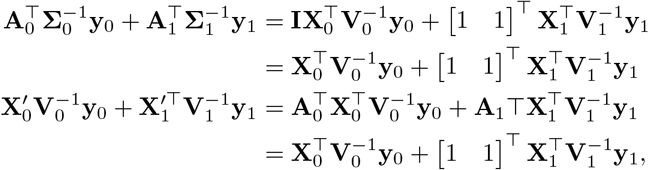

establishing the equivalence between (7) and (8), (9). In particular, the effect estimation variance-covariance matrices from the two approaches are the same. Note that the relative effective sample size for an effect estimate is given by the ratio of the estimation variance from the singletons to that from the augmented sample. Thus, it suffices to consider sample size gain in the meta-analysis approach.

#### 8.5.1 Imputation from sibling pairs

We give the theoretical effective sample size gain for direct effect estimate 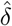 from adding *n*_1_ singletons to a sample of *n*_0_ sibling pairs. We consider a model in (1) with the parameter vector [*δ α*]^⊤^. From standard univariate GWAS on the *n*_1_ singletons, we have

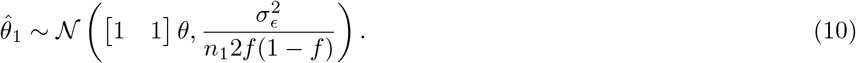

Let *r* be the sibling phenotypic correlation and *v* be the proportion of parental genotypic variance attributable to the imputation. Then, under random-mating, family-based GWAS yields[8]:

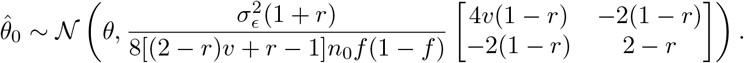

Following from (7), we have

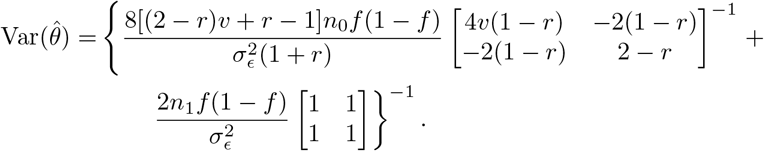

If imputation is done using phased IBD data, *v* = 3/4; if unphased data is used, *v* = 3/4 − *f*(1 − *f*)/8. Consider the special case where there is zero sibling phenotypic correlation, and that phased IBD data is used. Then

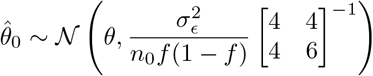

with

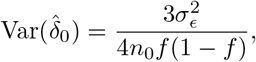

and

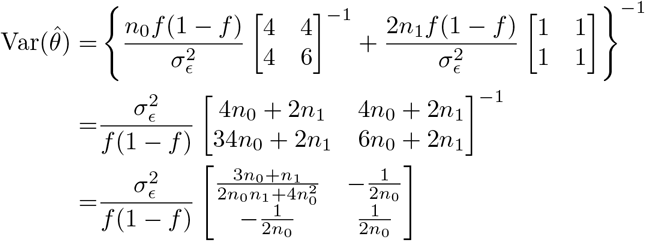

with

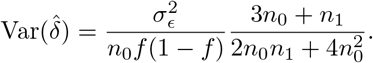

Therefore, the relative effective sample size is given by

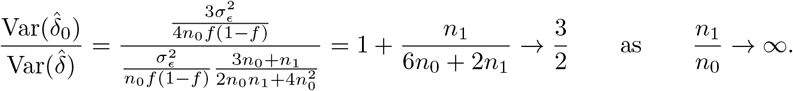

For other cases, the theoretical gain can be numerically computed for known *n*_0_ and *n*_1_.

To validate the theoretical derivation, we simulate datasets of 1500 sibling pairs with fixed allele frequency and varying sibling phenotypic correlations and compare the observed relative effective sample size to the theoretical values. From Figure 7a, we see that the observed values lie almost perfectly on the theoretical trend.

**Figure 7:**
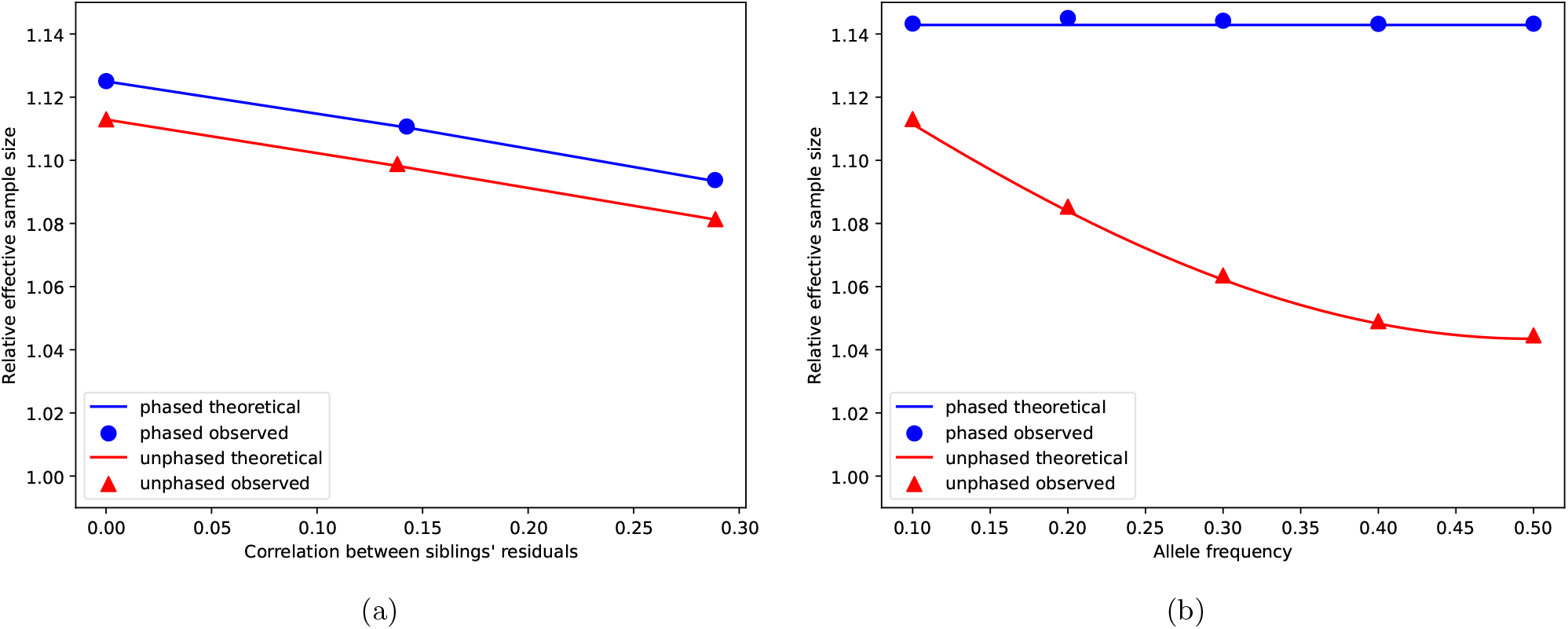
Observed and theoretical relative effective sample sizes. *Note*. We simulate 3,000 fathers’ and 3,000 mothers’ genotypes of 1,000 SNPs from independent binomials with 0.5 allele frequency. We then simulate meiosis and produce 3,000 sibling pairs while recording the IBD sharing information (to eliminate the influence of LD structure, we restrict the size of blocks without recombination to 1). Direct SNP effects are simulated from 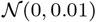. (a) We randomly remove genotypes of one sibling from 1,500 sibling pairs, resulting in *n*_0_=1,500 genotyped sibling pairs and *n*_1_=1,500 singletons. Parental genotypes of the *n*_0_ sibling pairs are imputed with and without phasing information, and those of the *n*_1_ individuals are linearly imputed. Then we perform two family-based GWAS analyses using *n*_0_ sibling pairs and *n*_0_ sibling pairs plus *n*_1_ individuals respectively, using Model 1. Observed relative effective sample sizes (circles and triangles are calculated by taking the ratio of direct effect estimation variance from the *n*_0_ sibling pairs to the estimation variance from the combined sample. The theoretical values are computed using expressions in section 8.5.1. (b) We randomly remove both parents’ genotypes of 1,500 offsprings, and fathers’ genotypes for the remaining 1,500 offsprings, resulting in *n*_0_=1,500 parent-offspring pairs and *n*_1_=1,500 singletons. Paternal genotypes of the *n*_0_ parent-offspring pairs are imputed with and without phasing information, and both parents’ genotypes of the *n*_1_ individuals are linearly imputed. Direct SNP effects are simulated from 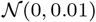. Then we perform two family-based GWAS analyses using *n*_0_ sibling pairs and *n*_0_ sibling pairs plus *n*_1_ individuals respectively, modeling direct effect and paternal and maternal NTCs. Observed relative effective sample sizes (circles and triangles are calculated by taking the ratio of direct effect estimation variance from the *n*_0_ parent-offspring pairs to the estimation variance from the combined sample. The theoretical values are computed using expressions in section 8.5.2.

#### 8.5.2 Imputation from parent-offspring pairs

We give the theoretical effective sample size gain for direct effect estimate 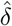 from adding *n*_1_ singletons to a sample of *n*_0_ parent-offspring pairs. Suppose mothers’ genotypes are observed fathers’ genotypes are imputed using phased data.

##### Modeling paternal and maternal NTCs

First consider a model in (2) with the parameter vector [*δ α_p_ α_m_*]^⊤^. From standard univariate GWAS on the *n*_1_ singletons, we have

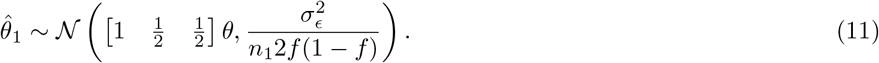

By section 5.3 of the supplementary note in [8], we have

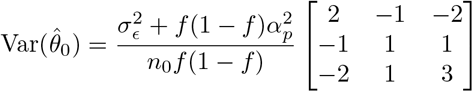

with

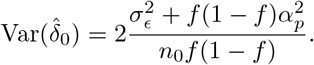

Again by (7),

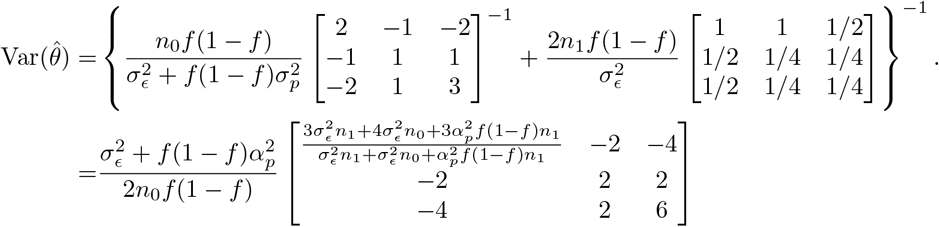

with

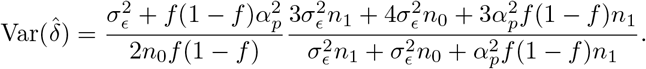

Then the relative effective sample size is given by

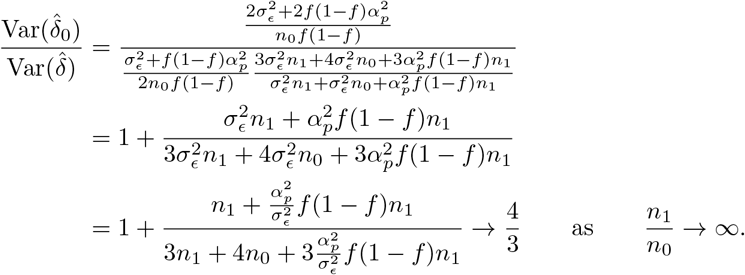

If 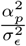 is close to zero,

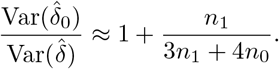

##### Modeling average NTCs

Now assume *α_p_* = *α_m_* and consider (1) as the true data generating model (see Appendix A for a discussion about violation of this assumption). From section 5.3 of the supplementary note in [8], we have

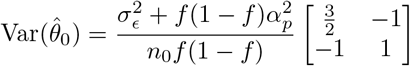

with

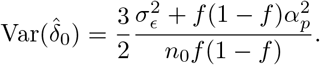

Similarly we have (10) from standard univariate GWAS. Again by (7),

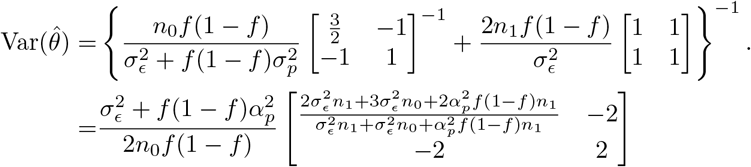

with

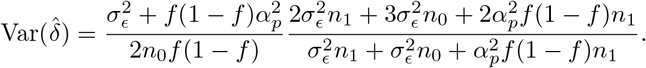

Then

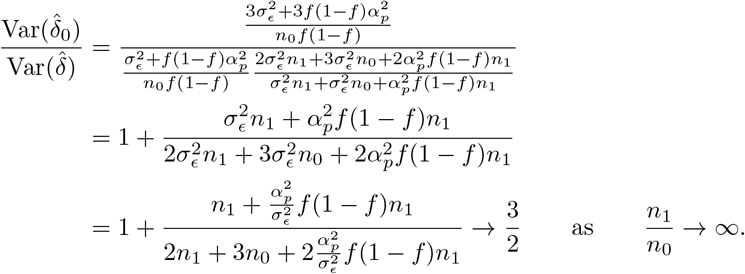

Note that in real-life datasets, 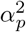 is usually trivial relative to 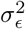. Thus we will have

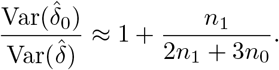

In the case where genotype data is unphased, the effective sample size gain is dependent on the allele frequency and can be computed numerically. We conduct simulations similar to the previous section, and the results are shown in Figure 7b.

### 8.6 Properties of the robust estimator

In this section, we give a derivation of the consistency and asymptotic unbiasedness of the the robust estimator in an island model of population structure. We proceed by first proving the claim for each group in table 2. For ‘paternal NT’, ‘maternal NT’, and ‘both NT’, we consider a single proband from each family, while for ‘one NT’, we consider sibling pairs. *n*_pat_, *n*_mat_, *n*_both_ and *n*_one_ are the numbers of families in the four groups, and families are assumed to be independent. The fully observed and imputed design matrices are given by **X** and 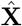 with the corresponding subscripts. We denote the parameter vectors for the ‘paternal NT’ and ‘maternal NT’ groups are *θ* = [*δ α_p_ α_m_*], and the parameter vectors for the ‘both NT’ are used to prove the claimed properties; and we give the sampling variances of the resulting estimators in each group.

We start by considering ‘both NT’, that is, the situation where both non-transmitted parental alleles are known. As all four parental alleles have been observed, the sum and imputed sum of the parental genotypes are the same: 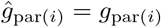, and the true and imputed design matrices coincide: 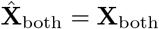, where 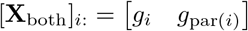. Denote the effect estimate by 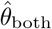. Then

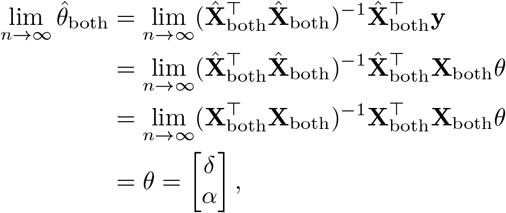

and specifically, the direct effect estimate 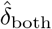 is consistent and asymptotically unbiased.

Next we look at ‘maternal NT’, the scenario where the non-transmitted maternal allele is observed and but the paternal one is not (‘Maternal NT’ in table 2). In the opposite situation ‘paternal NT’, the claim will follow by symmetry. In this case, the mother’s genotypes are completely determined: 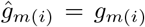. Then the imputed and complete design matrices are 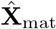 and **X**_mat_ respectively, with 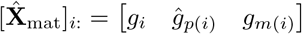 and 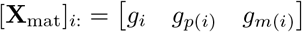. For the corresponding effect estimate 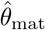 with paternal non-transmitted alleles imputed, we follow the supplementary note in Young et al. [8] and obtain (with a minor correction)

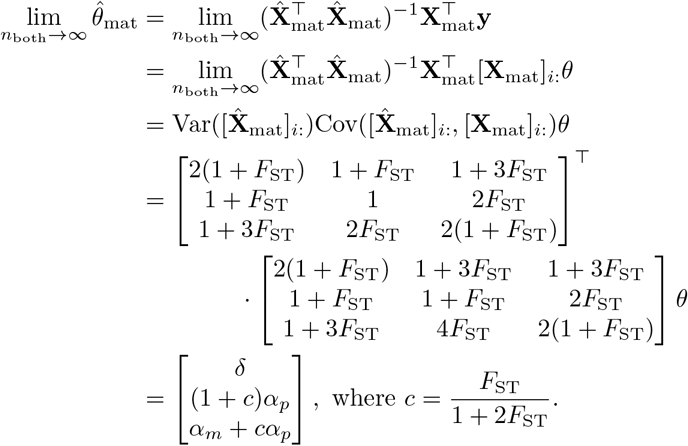

Therefore, the direct effect estimate 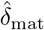 is consistent and asymptotically unbiased, although the non-transmitted coefficient estimates are biased. Similarly for ‘paternal NT’, we have

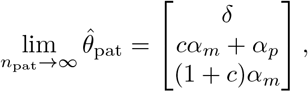

establishing consistency and asymptotic unbiasedness of 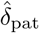.

Young et al. [8] show that the direct effect estimate 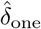 from sibling pairs in IBD1 (in fact, all possible cases in group ‘one NT’ in table 2 collapse to a sibling pair in IBD1) is also consistent and asymptotically unbiased:

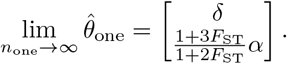

As is shown in (8.5), the robust estimator, which analyze the combined sample of the four groups in table 2, is equivalent to meta-analyzing 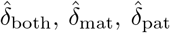 and 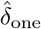. As these four estimators are consistent and asymptotically unbiased, so is the robust estimator.

**Table 5:** Limiting statistics for the four groups in Table 2.

| Group | $\text{Var}([\hat{\mathbf{X}}]_{i:})$ | $\text{Cov}([\hat{\mathbf{X}}]_{i:}, [\mathbf{X}]_{i:})$ |
| --- | --- | --- |
| Maternal NT | $f(1-f) \begin{bmatrix} 2(1+F_{\text{ST}}) & 1+F_{\text{ST}} & 1+3F_{\text{ST}} \\ 1+F_{\text{ST}} & 1 & 2F_{\text{ST}} \\ 1+3F_{\text{ST}} & 2F_{\text{ST}} & 2(1+F_{\text{ST}}) \end{bmatrix}$ | $f(1-f) \begin{bmatrix} 2(1+F_{\text{ST}}) & 1+3F_{\text{ST}} & 1+3F_{\text{ST}} \\ 1+F_{\text{ST}} & 1+F_{\text{ST}} & 2F_{\text{ST}} \\ 1+3F_{\text{ST}} & 4F_{\text{ST}} & 2(1+F_{\text{ST}}) \end{bmatrix}$ |
| Paternal NT | $f(1-f) \begin{bmatrix} 2(1+F_{\text{ST}}) & 1+3F_{\text{ST}} & 1+F_{\text{ST}} \\ 1+3F_{\text{ST}} & 2(1+F_{\text{ST}}) & 2F_{\text{ST}} \\ 1+F_{\text{ST}} & 2F_{\text{ST}} & 1 \end{bmatrix}$ | $f(1-f) \begin{bmatrix} 2(1+F_{\text{ST}}) & 1+3F_{\text{ST}} & 1+3F_{\text{ST}} \\ 1+3F_{\text{ST}} & 2(1+F_{\text{ST}}) & 4F_{\text{ST}} \\ 1+F_{\text{ST}} & 2F_{\text{ST}} & 1+F_{\text{ST}} \end{bmatrix}$ |
| Both NT | $f(1-f) \begin{bmatrix} 2(1+F_{\text{ST}}) & 2(1+3F_{\text{ST}}) \\ 2(1+3F_{\text{ST}}) & 4F_{\text{ST}} \end{bmatrix}$ | $f(1-f) \begin{bmatrix} 2(1+F_{\text{ST}}) & 2(1+3F_{\text{ST}}) \\ 2(1+3F_{\text{ST}}) & 4F_{\text{ST}} \end{bmatrix}$ |
| One NT | $\begin{bmatrix} (2-r) + (2-3r)F_{\text{ST}} & 2(1-r)(1+2F_{\text{ST}}) \\ 2(1-r)(1+2F_{\text{ST}}) & 3(1-r)(1+2F_{\text{ST}}) \end{bmatrix}$ | $f(1-f) \begin{bmatrix} (2-r) + (2-3r)F_{\text{ST}} & 2(1-r)(1+3F_{\text{ST}}) \\ 2(1-r)(1+2F_{\text{ST}}) & 3(1-r)(1+3F_{\text{ST}}) \end{bmatrix}$ |
| Group | $\text{Var}(\hat{\theta})$ | $\text{Var}(\hat{\delta})$ |
| Maternal NT | $\frac{\sigma_{\epsilon}^2}{2n_{\text{pat}}f(1-f)(1-F_{\text{ST}})} \begin{bmatrix} 2 & -2 & -1 \\ -2 & \frac{5F_{\text{ST}}+3}{2F_{\text{ST}}+1} & \frac{F_{\text{ST}}+1}{2F_{\text{ST}}+1} \\ -1 & \frac{F_{\text{ST}}+1}{2F_{\text{ST}}+1} & \frac{F_{\text{ST}}+1}{2F_{\text{ST}}+1} \end{bmatrix}$ | $\frac{\sigma_{\epsilon}^2}{n_{\text{pat}}f(1-f)(1-F_{\text{ST}})}$ |
| Paternal NT | $\frac{\sigma_{\epsilon}^2}{2n_{\text{mat}}f(1-f)(1-F_{\text{ST}})} \begin{bmatrix} 2 & -1 & -2 \\ -1 & \frac{F_{\text{ST}}+1}{2F_{\text{ST}}+1} & \frac{F_{\text{ST}}+1}{2F_{\text{ST}}+1} \\ -2 & \frac{F_{\text{ST}}+1}{2F_{\text{ST}}+1} & \frac{5F_{\text{ST}}+3}{2F_{\text{ST}}+1} \end{bmatrix}$ | $\frac{\sigma_{\epsilon}^2}{n_{\text{mat}}f(1-f)(1-F_{\text{ST}})}$ |
| Both NT | $\frac{\sigma_{\epsilon}^2}{n_{\text{both}}f(1-f)(1-F_{\text{ST}})} \begin{bmatrix} 2 & -1 \\ -1 & \frac{F_{\text{ST}}-1}{3F_{\text{ST}}+1} \end{bmatrix}$ | $\frac{\sigma_{\epsilon}^2}{2n_{\text{both}}f(1-f)(1-F_{\text{ST}})}$ |
| One NT | $\frac{(1+r)\sigma_{\epsilon}^2}{2n_{\text{one}}(2+r)(1-F_{\text{ST}})(1+2F_{\text{ST}})f(1-f)} \begin{bmatrix} 3(1-r)(1+2F_{\text{ST}}) & -2(1-r)(1+2F_{\text{ST}}) \\ -2(1-r)(1+2F_{\text{ST}}) & (2-r) + (2-3r)F_{\text{ST}} \end{bmatrix}$ | $\frac{3(1-r^2)\sigma_{\epsilon}^2}{2n_{\text{one}}(2+r)f(1-f)(1-F_{\text{ST}})}$ |

## A Modeling average NTC with imputation from parent-offspring pairs

One somewhat surprising finding is that assuming *α_p_* = *α_m_* can produce a more precise estimate of *δ* when analyzing parent-offspring pairs with imputation of the missing parent’s genotype. In the case of imputing the missing father’s genotype given the mother and offspring’s genotype, the portion of variance of the proband’s genotype *g_ij_* that is uncorrelated with 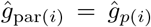 is larger than the portion uncorrelated with both 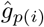 and *g*_*m*(*i*)_, and thus we have more information for estimation of *δ*, at the cost of bias when *α_p_* ≠ *α_m_*. We establish this result in this section.

Suppose we have *n* mother-offspring pairs with observed and phased genotypes, *g_i_* and *g*_*m*(*i*)_. Then the father’s genotype *g*_*p*(*i*)_ can be imputed the same way as sibling pairs in IBD2:

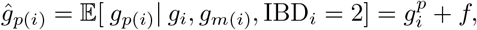

where 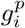 is the offspring allele inherited from the father. We model the phenotype using (1), i.e., average NTC is modeled instead of paternal the maternal NTCs modeled separately. So we have 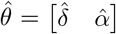 and the imputed design matrix 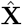 with 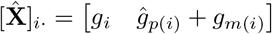.

If the phenotype is generated by (2), we have the parameter vector *θ* = [*δ α_p_ α_m_*], and the complete design matrix **X** with [**X**]_*i*·_ = [*g_i_ g*_*p*(*i*)_ *g*_*m*(*i*)_]. Then

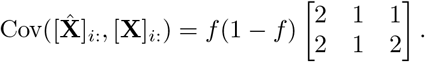

Therefore,

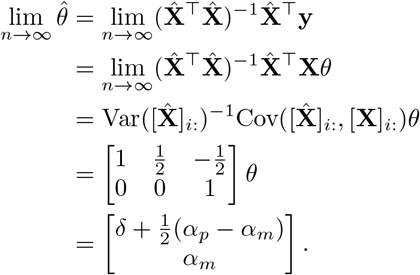

If *α_p_* ≠ *α_m_*, the OLS estimate 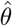 is inconsistent and asymptotically biased.

Assume instead *α* = *α_p_* = *α_m_*, then the phenotype generating model is equivalent to (1). Then *θ* = [*δ α*] and 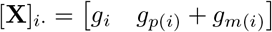. Thus we have

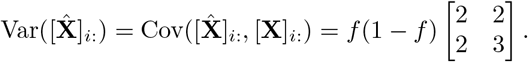

Then,

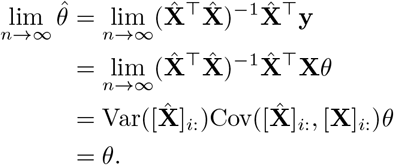

i.e., the OLS estimate 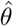 is consistent. Also,

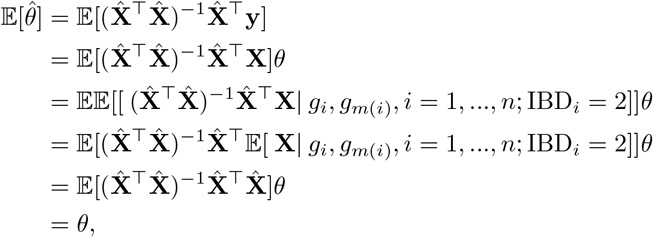

i.e., 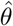 is unbiased. By the supplementary note in [8], we have

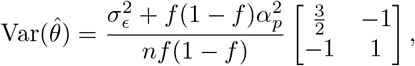

with

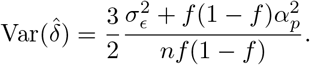

For *θ_c_* = [*δ α_p_ α_m_*], we have

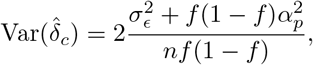

and 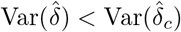.

